# Neural Signatures of Automatic Auditory Regularity Detection Reflect Individual Differences in Explicit Short-Term Memory

**DOI:** 10.64898/2026.01.09.698639

**Authors:** Mingyue Hu, Maria Chait

## Abstract

Perception is shaped by the statistical structure of the environment, reflecting the brain’s capacity to detect and exploit regularities in sensory input. This process requires the maintenance of contextual information over time, yet the nature of the memory mechanism supporting automatic structure learning remains unclear. Here, we ask whether auditory regularity processing relies on a dedicated sensory buffer or instead draws on domain-general mnemonic resources. We related neural indices of regularity processing measured with EEG during passive listening to behavioural performance on an explicit auditory short-term memory task. Human participants (N=30; both sexes) passively listened to regularly repeating (cycles of 5.5 seconds) or random tone sequences, while sustained and tone-locked neural responses were extracted as complementary markers of predictability tracking and prediction-error signalling. Individual differences in explicit memory performance, quantified using a delayed match to sample task, systematically predicted both neural measures: high performers showed enhanced sustained responses and attenuated tone-evoked responses (most pronounced in the N2 time window; 250–400 ms post-onset) to regular sequences, whereas low performers showed no reliable modulation by sequence structure. These findings demonstrate that the memory processes engaged automatically during auditory pattern analysis are not encapsulated, but instead draw on shared mnemonic resources, providing a link between predictive perception and individual variability in sensory memory capacity.

## Introduction

Perception is fundamentally shaped by the statistical structure of the environment, exhibiting a pronounced sensitivity to the regularities embedded in sensory input (1–18). Within contemporary theoretical frameworks, this capacity is understood as a core component of predictive inference, whereby the brain constructs internal generative models that capture probabilistic regularities in sensory input (9). These models are used to generate expectations about forthcoming events, and discrepancies between predicted and observed input are evaluated to update the model and minimize prediction error over time (2,8). By continuously optimizing these internal representations, the brain can allocate cognitive and neural resources efficiently, supporting adaptive perception and action in a dynamically changing environment (3,10–15).

Converging evidence indicates that the neural mechanisms supporting the extraction and tracking of auditory statistical regularities can be investigated using sustained activity measured with M/EEG. In the canonical paradigm (e.g., (9)), listeners passively hear tone sequences that are either periodically repeating (REG) or randomly ordered (RND). REG sequences elicit a monotonic increase in sustained neural activity that asymptotes as the underlying regularity is inferred, consistent with the formation of a dynamic neural representation of sequence structure (25–28). For sequences with pattern lengths of up to ten tones, the latency at which REG and RND responses diverge—approximately three tones following completion of the first cycle—closely matches predictions derived from an ideal observer model of statistical learning (9,16–18). This temporal correspondence indicates statistical efficiency in the neural encoding of sequential structure, even in the absence of task relevance or explicit behavioural engagement (9). Source localization studies implicate a distributed network including auditory cortex, inferior frontal gyrus (IFG), and hippocampus as key substrates supporting this effect (9,19,20). A growing body of work further shows that the sustained response scales systematically with the precision of sequential auditory input and is observed for both deterministic and stochastic regularities, providing a direct neural index of online statistical inference and adaptive tracking of environmental structure (9,14,21–25).

Using sequences in which tones were temporally separated by silent gaps—allowing dissociation of sustained and tone-locked responses—Hu et al. (24) demonstrated that modulation of sustained activity occurs concurrently with opposing effects at the level of individual tones. Specifically, the establishment of regularity is associated with increased sustained responses alongside attenuated tone-evoked responses, consistent with reduced prediction error. These complementary effects indicate parallel neural processes that jointly shape the representation of unfolding auditory structure and align with the formation of a top-down predictive model of the sequence.

The maintenance of contextual information necessary for determining predictability requires a memory store. Several lines of work suggest that this store may be a source of individual variability in sensitivity to regularity, as shown by studies in which the sustained response is systematically modulated by manipulations that increase mnemonic load. These manipulations include lengthening the tones comprising a pattern (19), increasing the number of tones per regularity cycle (9,16), and extending cycle duration by inserting silent gaps between tones (24). Across these conditions, sustained activity is attenuated and shows increased inter-participant variability, consistent with the hypothesis that the system approaches the limits of automatic auditory memory capacity.

Further support for this interpretation comes from lifespan research. Herrmann et al. (26) examined age-related differences in neural responses to auditory patterns and reported that sustained activity to regular sequences was significantly reduced in older relative to younger adults. This dissociation suggests an impairment in the ability to integrate and maintain sequential information over time, potentially reflecting age-related declines in auditory sensory memory.

Because predictive models rely critically on memory capacity to retain and integrate sequential information over extended timescales, characterizing the nature of this memory buffer is essential for understanding automatic structure learning, inter-individual variability, and vulnerability to impairment. A central unresolved question is whether this buffer constitutes a dedicated, encapsulated sensory memory system, or whether it draws on domain-general mnemonic resources shared with higher-level cognitive functions.

Here, we directly tested the hypothesis that individual differences in explicit auditory short-term memory capacity are linked to the efficiency with which listeners automatically track auditory statistical structure by relating automatic neural indices of regularity processing—measured using EEG during passive exposure to structured sound sequences—to behavioural performance on an explicit auditory short-term memory task. The auditory stimulus used in the passive listening paradigm consisted of a REG sequence comprising a 10-tone pattern. To maximally tax auditory memory resources, tones were separated by 500-ms silent gaps, yielding a cycle duration of 5.5 s. Previous work has shown that this temporal regime produces substantial behavioural variability in pattern detection, consistent with limits on automatic memory integration (24). EEG was recorded while participants listened passively to these sequences. We extracted both sustained activity and tone-locked responses to quantify complementary neural signatures of regularity processing. Explicit memory was assessed, post EEG session, using a variant of the two-interval comparison paradigm commonly employed to probe memory for tone sequences (27,28) (Fig. 2). This approach allows us to assess the extent to which automatic neural tracking of auditory structure is constrained by, and predictive of, explicit task-dependent sensory memory capacity.

## Methods

### Participants

Thirty-four naïve paid participants with normal pure-tone thresholds (≤ 20 dB) in standard audiometric frequencies (0.25–8 kHz) participated in the study. Data from two participants were discarded due to excessive noise, and data from another two were discarded due to a trigger malfunction resulting in data loss. Hence, data from 30 participants (19 female; average age, 24.7 ± 4.63) are reported below. All were fluent in English, had normal or corrected-to-normal vision. The research ethics committee of University College London approved all experimental procedures described in this study, and written informed consent was obtained from each participant.

### Procedure

The experiment consisted of two main stages: Participants first completed an **EEG** measurement in which they listened to tone patterns whilst watching a silent movie. It was followed by a brief **behavioural assay** where auditory sensory memory was measured using a tone sequence comparison task. Participants were only informed of this task after the EEG session has concluded.

The experiment was implemented in Psychophysics Toolbox in MATLAB (29) and conducted in a soundproof booth. EEG signals were recorded with a Biosemi system (Biosemi Active Two AD-box ADC-17, Biosemi, Netherlands) with 64 Ag-AgCl electrodes at a 2048 Hz sampling rate and subsequently downsampled to 256 Hz. The recording was restarted for each block. Auditory stimuli were delivered binaurally via tube earphones (EARTONE 3A 10 Ω; Etymotic Research) inserted into the ear canal. The loudness was adjusted to each participant’s comfort. The measurement lasted a total of 40 minutes.

**The EEG session** began with an auditory functional localizer block, lasting about 3 minutes. This block contained a randomized sequence of 180-200 pure tones (1000 Hz frequency, 150 ms duration), each followed by a random interstimulus interval (ISI) between 700 and 1500 ms. This process served as a control to ensure a reasonable signal-to-noise ratio. In the main experiment, participants passively listened to randomly presented tone sequences (see below) with an inter-stimulus interval (ISI) of 3000-4500 ms, while watching a silent movie of their choice. Unaware of the auditory stimuli’s nature, participants were encouraged to focus on the movie. The session was divided into five 8-minute blocks, with short breaks allowed between blocks while participants remained still.

Stimuli (Figure 1A) were sequences of 50-ms tone-pips, each gated on and off with 5-ms raised cosine ramps. Frequencies were selected from a pool of 20 values equally spaced on a logarithmic scale from 222 to 2000 Hz (12% steps). Two sequence types were created: **REG** sequences were generated by randomly selecting 10 frequencies from the pool without replacement. These were arranged in random order to form a pattern, which was then repeated three times. New REG patterns were generated for each trial. **RND** sequences also consisted of 10 frequencies, similarly, selected anew for each trial and presented in random order. All sequences contained 30 tone-pips, presented with a 500 ms inter-tone silent gap (1.8 Hz rate; 5500ms REG cycle duration; 16.5 sec overall sequence duration). Stimulus delivery was in blocks. Each block consisted of 10 REG and 10 RND trials in a randomized order.

**Figure 1:**
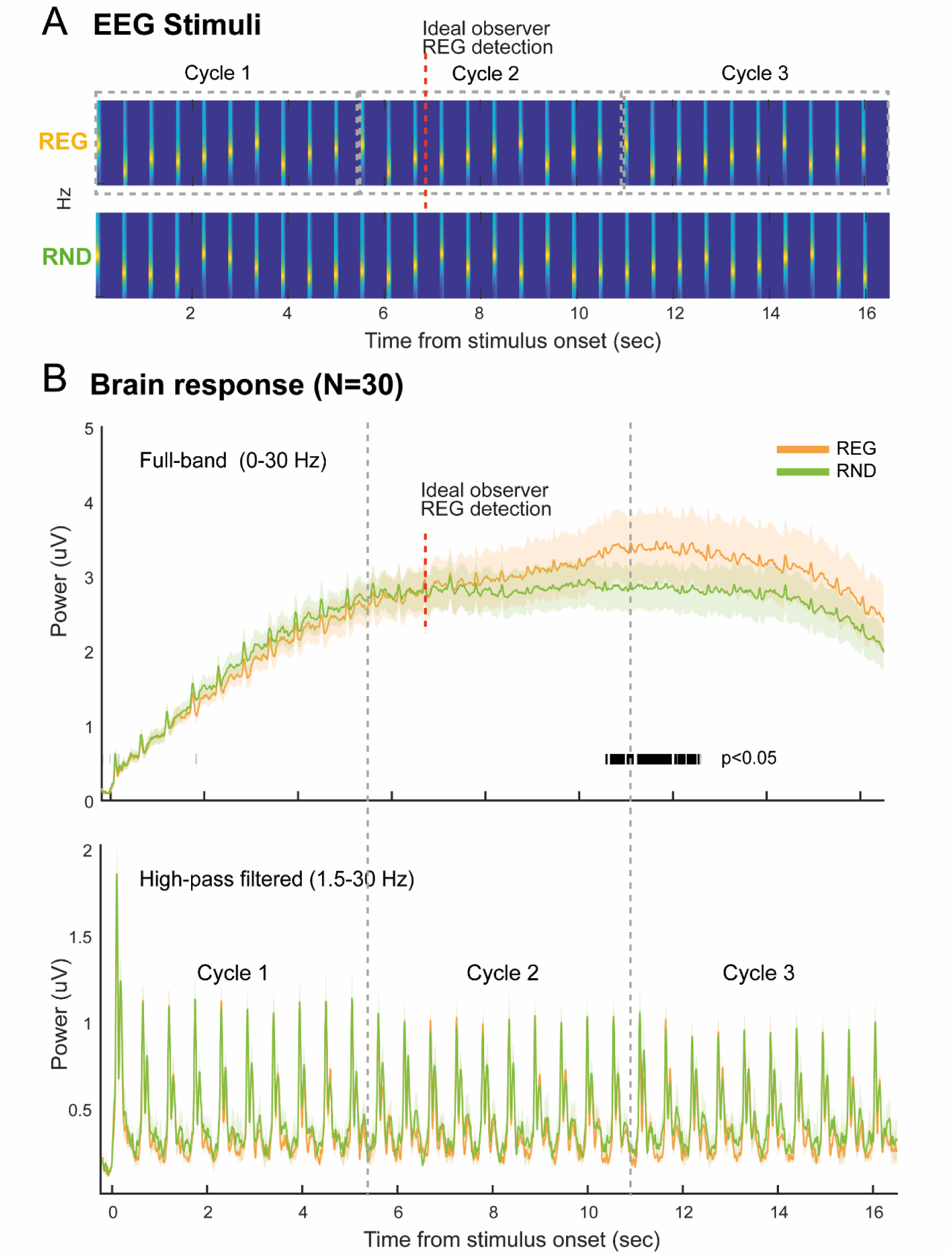
EEG stimuli and main responses. **(A)** Spectrogram of example REG and RND sequences used in the EEG experiment. Signals consisted of thirty 50 ms tones (3 regularity cycles in REG sequences; 16.5 s long), with 500 ms silent gaps between tones. Naïve participants listened to the sound sequences passively while watching a movie of their choice. We hypothesized that if the brain monitors the transition probabilities between tones in the unfolding sequence, responses to REG and RND should be differentiated during the second cycle (when the pattern begins to repeat). Ideal observer-based estimates (e.g. (9); (16)) suggest an ideal observer requires roughly 3 tones (marked by the dashed red line) after the onset of cycle 2 to distinguish between REG and RND. **(B)** EEG response evoked by REG and RND sequence. The top panel displays group RMS across a frequency range of 0-30 Hz. The two traces represent the average power during the presentation of REG vs RND tone sequences. The shaded areas denote the standard error. The black bars below the graph indicate statistically significant (p<0.05) differences between the two conditions. The bottom graph shows group RMS in a narrower frequency band of 1.5-30 Hz which emphasizes the tone-evoked activity (1.8 Hz).

**The behavioural memory task** used - Tone Pattern Comparison Task (TP-COMP) - was similar to that reported in (28): The stimuli consisted of two 500 ms tone-pip sequences separated by a 2000 ms silent gap (see Figure 2A). The sequences comprised ten 50 ms tone-pips drawn from the same pool used for the EEG stimuli above. Different patterns were drawn on each trial. The two sound sequences before and after the gap were matched on 50 % of the trials (‘same’ trials) and differed in the other trials (‘different’ trials). The sequences in the ‘different’ trials were created by switching the positions of 3 of the 10 tones. The positions of the shuffled tones were randomly chosen on each trial, excluding the first and last tones, to avoid primacy and recency effects. The instructions were to listen carefully to the sound sequences and press one of two keyboard buttons to indicate whether the two-tone sequences were the same or different (“S” for same and “D” for different). Participants then completed 32 trials. Feedback was provided after each trial. The correct response rate was used to quantify performance. Before proceeding to the main task, subjects were given a short practice (10 trials). Stimuli were delivered over Sennheiser HD558 headphones via a UA-33 sound card. The testing took place in the same booth as the EEG experiment. The task, including instructions, took approximately 5 min to complete.

**Figure 2:**
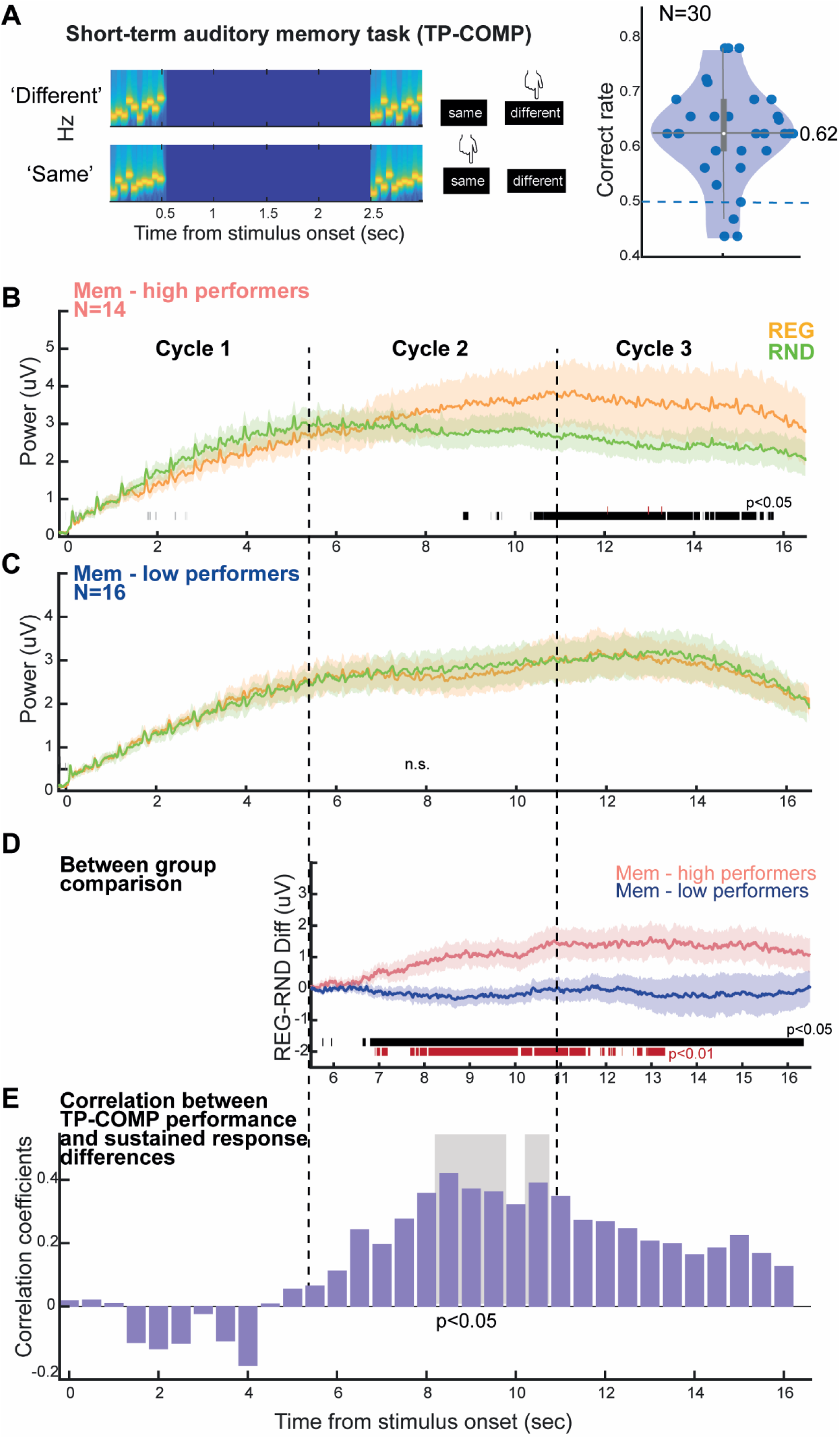
Sequence evoked EEG responses grouped based on TP-COMP performance. **(A)**: Left: Examples of the stimuli used in the Tone Pattern Comparison Task (TP-COMP). Participants listened to two 500 ms tone sequences (10 random tone pips), separated by a 2-second silent interval. They were then asked to indicate whether the two sequences were the same or different. In ’different’ trials, the positions of three tones (but never the first or last tone) were altered. Right: Task performance, expressed as % correct. Chance level (0.5) is marked with a dashed line. The white circle at 0.62 indicates the median. **(B)** EEG traces (Group RMS, 0-30 Hz) for participants who scored above the median in the TP-COMP task (N=14). Horizontal bars below the traces indicate intervals of significant difference between conditions. Any significant differences observed in the baseline period or during cycle 1 (where REG and RND are identical and differences are therefore not expected) are indicated in grey and used as a threshold for correcting responses during cycle 2 and 3 (see methods). **(C)** EEG traces (Group RMS) for participants who scored below or equal to the median on the TP-COMP task (N=16). **(D)** Direct comparison between memory performance groups. The plot compares the EEG response differences of REG and RND between memory high performers and memory low performers. **(E)** Spearman correlation between sustained response differences (REG-RND), binned in 0.5 sec steps, and TP-COMP performance. Purple bars represent the Spearman correlation coefficient at each bin. Grey shaded areas mark the time intervals where a significant correlation (p < 0.05; FWE uncorrected) was observed.

## EEG Data analysis

### Detrending

Before preprocessing, robust detrending was performed using a 10th order polynomial fit (30), with the fitted values subsequently subtracted from the original continuous signal. This was done to reduce common drifts in EEG recording, especially since the stimulus used in this study is particularly slow. This approach (as opposed to high-pass filtering) helps preserve the slow dynamics (sustained response) that are phase-locked to the stimulus and are of interest in this study.

### EEG Data Preprocessing

All pre-processing and time domain analyses were conducted using the fieldtrip toolbox (http://www.fieldtriptoolbox.org/, (31). Low-pass filtering was applied at 30 Hz (all filtering in this study was performed using a two-pass, Butterworth filter with zero phase shift). To analyse time domain data, we identified the 10 most responsive channels for each subject. This was done by combining tone responses collapsed from all conditions and identifying the N1 component (80-120 ms) of the onset response (32,33). For each subject, the 10 most strongly activated channels at the peak of auditory N1 (5 most positive, 5 most negative) were selected to best represent auditory activity for all subsequent time-domain analyses. This procedure served the dual purpose of enhancing the relevant response components and compensating for any channel misalignment between subjects.

### Sequence Evoked Response

First, we focused on analysing responses to the sequence, with special attention to low frequency activity as a potential marker of predictability tracking (9,16,24). We avoided using a high-pass filter. The data were segmented into epochs of 16.5 seconds, starting from 200 ms prior to onset until the end of each trial. These epochs were then baselined to the pre-onset interval (200 ms) and averaged for evoked response analysis.

To minimize low frequency drift artifacts and enhance signal noise ratio that is locked to the trial, we applied denoising source separation (DSS), following the method in (34), three components, which exhibited the highest reproducibility across trials, were identified and projected back into sensor space for each subject.

### Tone Evoked Response

A secondary analysis was focused on neural responses to individual tones in REG versus RND sequences. To identify activity linked to each tone-evoked response, which might be obscured by slow neural dynamics, the raw data were high-pass filtered at 1.5 Hz. The filtered data were then segmented into individual tone epochs, spanning from 50 ms before to 500 ms after tone onset. DSS was applied to the tone-evoked responses and the three most significant components were projected back to the sensor space for each participant. Responses from tones within each cycle were then averaged, producing three time series. This was done for each condition per subject. The time series were baselined based on the activity before the onset of the tone (50 ms).

### Statistical Analysis

The time domain data is summarized as root-mean square (RMS) across ten most responsive channels for each subject. RMS is a useful summary signal because it reflects the instantaneous power of the neural response, regardless of polarity. For illustration, we show the group-response (average of individual RMSs) and standard error across subjects. However, statistical analysis is always conducted across subjects. To assess differences between conditions (RND vs REG), we calculated RMS differences at each time point for each participant. We then applied a bootstrap resampling (35) with 1000 iterations to the entire epoch. We considered a significant difference if the proportion of bootstrap iterations that were either above or below zero exceeded 95% (i.e., p < 0.05). For time intervals where significant differences were not theoretically expected (e.g. during the baseline period, or during the first cycle where REG and RND are indistinguishable), any identified clusters were presumed to be due to noise (marked as grey in the figures below). The longest cluster identified in this way was used as a threshold for significance for the rest of the epoch, such that only clusters that exceed this duration are considered to be significant.

## Results

EEG responses to auditory sound sequences were examined in thirty participants and related to short-term memory abilities probed with a Tone Pattern Comparison Task (TP-COMP).

### Sequence-Evoked EEG Responses Suggest Regularity Extraction

The study explores brain responses to sound sequences, with a particular focus on the slow, sustained EEG response. The core question being investigated is whether the brain can automatically (outside of behavioural relevance) recognize slow REG patterns (cycle duration of 5500 ms). This is being studied by testing whether brain responses to REG patterns differ from matched random sequences (RND). The upper graph of Figure 1B presents the group EEG response (mean and standard error of individual RMSs) to RND and REG sequences. A typical onset response is observed, which is then followed by a heightened level of sustained activity. This ongoing activity is punctuated by clear fluctuations at 1.8 Hz, mirroring the frequency at which the tones were presented. This observation is consistent with past MEG work, which has reported a similar pattern of sustained neural activity for patterns presented at 4 Hz (24).

At the whole group level, we observed a subtle enhancement in this sustained neural activity evoked by REG sequences in comparison to RND ones, particularly evident from the third cycle onwards.

### Performance in the tone pattern comparison task exhibits significant variability

The Tone Pattern Comparison Task (TP-COMP(28); Figure 2A) used here is similar to previous methods for evaluating participants’ auditory short-term memory (36–38). Participants engage with this task by listening to pairs of tone sequences and determining whether each pair is same or different. Our analysis reveals significant variability in participant performance, as depicted in Figure 2A. The distribution of correct scores is consistent with a prior study (28), with mean and median around 62% but quite a large variability across participants, likely reflecting variance in short-term memory capacity.

To explore the relationship between task performance and brain activity, we divided participants into two groups based on a median split (“Mem – high”; “Mem – low”).

### TP-COMP performance is associated with differential EEG sustained responses to REG patterns

High-performing participants (Mem–high) exhibited a pronounced differentiation in sustained EEG responses to REG relative to RND sequences, expressed as an increase in power that emerged from the second cycle of REG presentation onward (Figure 2B). In contrast, low-performing participants (Mem–low) showed no reliable differentiation between EEG responses to REG and RND sequences (Figure 2C). As illustrated in Figure 2D, the interaction between group and condition (REG–RND) was significant, with effects emerging from the third tone of the second regularity cycle onward.

To further examine potential correlations between short-term memory performance and sustained response dynamics, we conducted an exploratory Spearman correlation analysis between the difference in sustained response (REG-RND) and the performance on TP-COMP. The correlations were conducted over 0.5 s intervals, sampling the entire epoch (Figure 2E). The analysis revealed significant effects (p<0.05; FEW uncorrected), mainly within the 7.5-11 sec time interval, which corresponds to cycle 2 of REG – the initial period over which REG is being discovered. The fact that the strongest correlation is observed during the second cycle suggests that this relationship is specifically linked to the **speed of regularity discovery**, which occurs during early exposure to the repeating pattern (cycle 2).

### Tone-evoked responses are modulated by context predictability

Following the analysis of sequence-evoked responses, we examined **phasic tone-evoked activity** (1.5–30 Hz) elicited by tones presented within REG and RND sequences. Figure 3 shows the group-averaged tone-locked responses, averaged separately across tones within each REG cycle and across the corresponding tones in the RND condition. The EEG response to individual tones was characterized by a sequence of distinct RMS peaks, as indicated in Figure 3.

**Figure 3:**
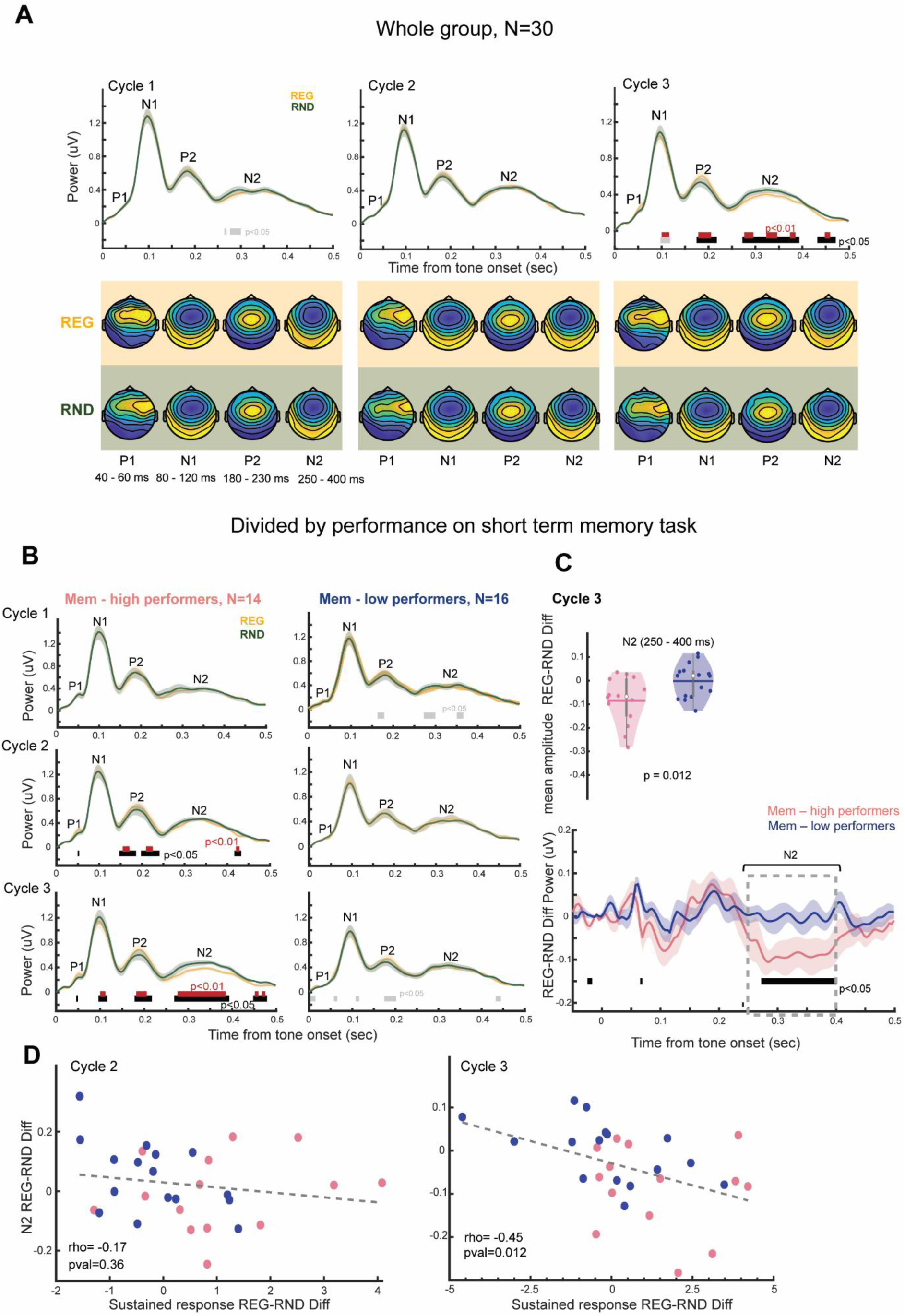
Tone-locked responses. **(A)** Tone evoked activity in cycle 1,2,3. Four distinct ERP components are visible, P1 (40-60ms), N1 (80-120ms), P2 (180-230ms), N2 (250-400ms). Topographical maps are provided below the EEG traces. Any differences between conditions during cycle 1 (indicated in light grey) are considered noise and used as a threshold of significance in cycle 2 and cycle 3 (see methods). Black (p<0.05) and red (p<0.01) horizontal lines mark intervals where a statistically significant difference is observed in cycles 2,3; Grey horizontal bars indicate intervals that did not pass the cluster threshold. **(B)** Tone-locked responses in TP-COMP high and low performers. Left column shows tone responses from high performers in TP-COMP (N=14) and the right shows those from low performers (N=16). Horizontal bars denote significant time intervals (as in Panel A). Grey bars indicate noise-attributed clusters (see methods). **(C)** Between group comparison during cycle 3 within the N2 time window. The bottom graph shows the tone response difference (REG-RND) for each group. Statistical comparison suggests a significant difference in the N2 time window (250-400ms) between groups. **(D)** Spearman correlation between mean amplitude of sustained response differences (REG-RND) and REG-RND differences during the N2 response (0.25-0.4s) in cycle 2 and 3.

Differences between conditions emerged during the third cycle. Specifically, tone-evoked responses were reduced for REG relative to RND sequences in the N1 and N2 time windows, whereas the opposite pattern was observed in the P2 window, with larger responses for REG than RND.

### TP-COMP performance Associated with Differential tone-evoked responses to REG patterns

In parallel with the sequence-evoked response analysis, we examined the relationship between tone-evoked neural responses and short-term memory performance. Figure 3B shows the averaged tone-evoked activity for each REG cycle and the corresponding time intervals in the RND condition. No significant effects were observed in the Mem–low group. In contrast, significant activity clusters were observed in the Mem–high group, as indicated in Figure 3B.

Figure 3C illustrates the interaction between participant group (high vs. low performers, defined by TP-COMP scores) and sequence type (REG vs. RND). Significant clusters were detected exclusively within the N2 time window (approximately 250–400 ms). To further characterize this effect, an independent-samples t-test was conducted on the mean amplitude of the N2 time window. This analysis confirmed a significantly reduced N2 amplitude in Mem–high participants compared with Mem–low participants [t(29) = −2.67, d = −0.98, p = 0.012]. Finally, we performed a Spearman correlation across all subjects, on the REG-RND difference in the tone-evoked response (N2 time window), and the REG-RND difference in the sustained response (as shown in Figure 1,2) during Cycles 2 and 3. This revealed a significant correlation between the N2 response and the sustained response (rho = -0.45, p = 0.012) during cycle 3 but not during cycle 2, consistent with the data in Figure 3B. The negative sign of the correlation is consistent with the opposite effects of regularity on the sustained response (REG>RND) and the tone evoked response (RND>REG).

### Memory task high performers exhibit enhanced early auditory responses

We next examined whether baseline auditory responses, collapsed across REG and RND tone-evoked activity during cycle 1, differed as a function of performance on the TP-COMP task (Figure 4). Analysis of tone-evoked responses, across both REG and RND conditions, revealed that participants with TP-COMP scores above the median exhibited significantly larger P1 and N1 amplitudes. This pattern suggests more efficient early sensory encoding of auditory input in these individuals. Notably, this enhancement was not observed in later evoked components typically associated with higher-order or integrative processing.

**Figure 4:**
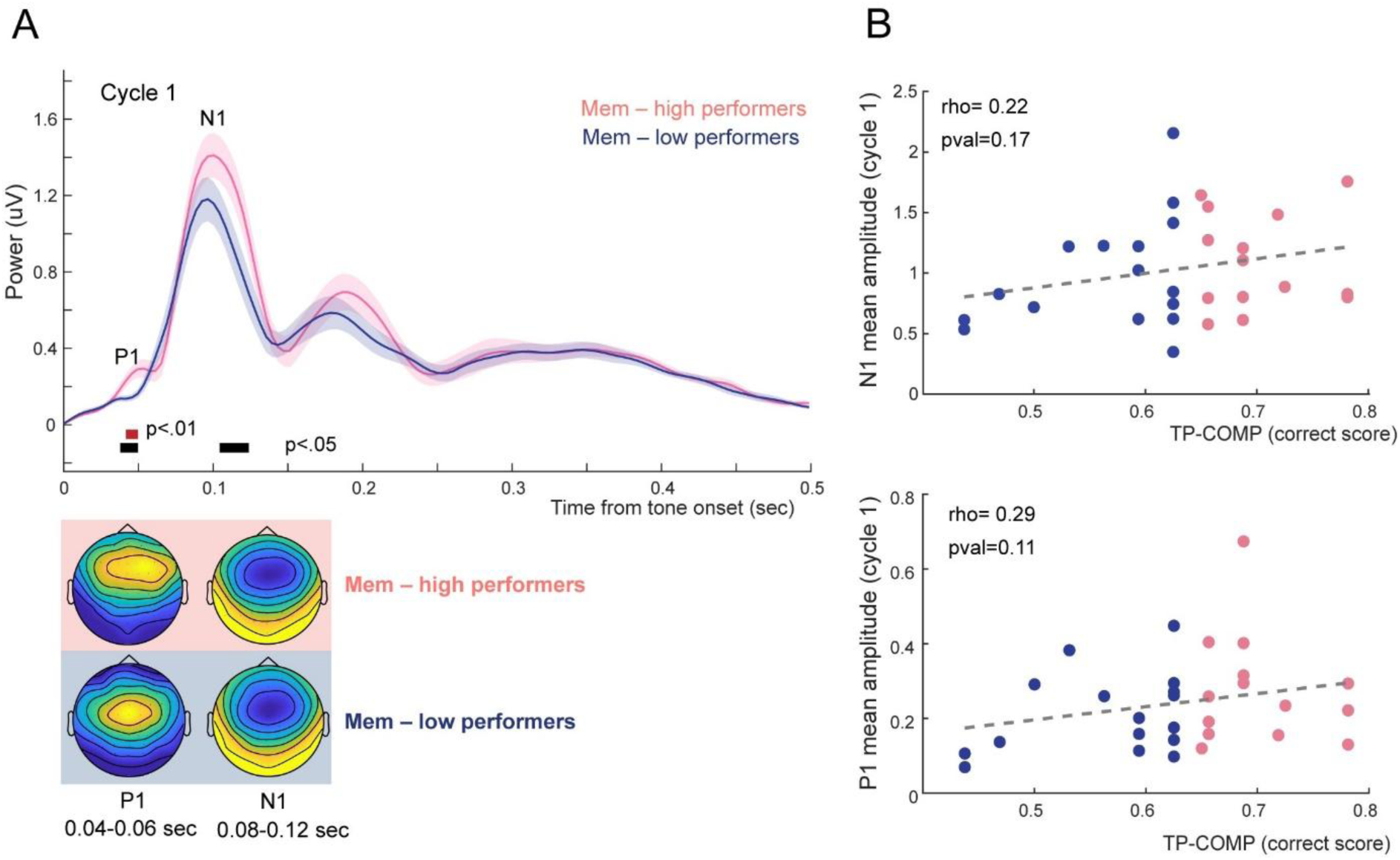
Tone-evoked activity in cycle 1. **(A)** Tone responses averaged across the first cycle collapsing across RND and REG. Statistically significant differences between groups were observed in the P1 (40-60ms) and N1 (80-120ms) time ranges. **(B)** Scatter plots for P1 and N1 response amplitudes against TP-COMP scores. In both cases the correlation (Spearman) is not significant.

To further assess the relationship between neural responses and behavioural performance, we conducted Spearman correlation analyses between TP-COMP scores and P1 and N1 amplitudes. As shown in Figure 4B, no significant correlations were observed for either N1 (ρ = 0.22, p = 0.17) or P1 (ρ = 0.29, p = 0.11).

## Discussion

The ability to discover predictable structure in the environment is a central tenet of predictive coding. Previous work has shown that neural mechanisms supporting the detection of regularity are underpinned by a distributed network encompassing auditory cortex, frontal regions, the basal ganglia, and the hippocampus (9,19,20,24). The dynamics of activity within this network are reflected in distinct components of M/EEG responses. Sustained low-frequency activity has been associated with tracking predictability (25) and typically exhibits larger responses for predictable compared with random patterns. In parallel, phasic event-related responses are thought to index prediction error processing and show reduced responses to tones embedded in predictable relative to random sources (24,39).

Together, these sustained and phasic responses are hypothesized to reflect the functionally distinct neuronal subpopulations proposed by predictive coding models (40–42): one population encodes conditional expectations about the causes of sensory input, while another encodes prediction errors. Their interaction enables the brain to both enhance the fidelity of expected sensory signals (43–45) and prioritize unexpected stimuli, which carry high informational value (46–48).

Here, we sought to understand how these processes relate to memory, given that memory is fundamental to predictive coding. We employed sequences composed of repeating patterns of 10 tones, with tones separated by 500 ms of silence, yielding pattern repetitions every 5.5 seconds. Behaviourally, participants show substantial variability in their ability to detect structure in these sequences, with a mean sensitivity (d′) of approximately 0.9 (24). We therefore expected these sequences to elicit considerable individual variability in EEG responses and selected them for the present investigation. Our results reveal that the memory resources engaged automatically during auditory pattern detection are not encapsulated, but rather shared with explicit short-term memory processes, providing a mechanistic link between memory, predictive coding, and individual differences in perception.

### Multiplexed representation of sequence predictability and event prediction error in the EEG signal

Consistent with Hu et al. (24), we observed evidence for sustained response differences between REG and RND sequences. Predictability in such sequences has often been modelled using ideal observer approaches based on variants of Prediction by Partial Matching (PPM), which estimate the probability of upcoming events from previously encountered sequences across a hierarchy of context lengths. This framework has successfully accounted for a wide range of behavioural and neural effects observed in human listeners (9,18,19,49–54). According to these models, under perfect memory conditions, the REG pattern should, in principle, become detectable after one full cycle plus three additional tones (approximately 7,150 ms here (9,18)). Inspection of the mean traces in Figure 1 suggests that a divergence between REG and RND responses emerges around this time; however, statistically reliable differences appeared substantially later and were relatively weak.

Analysis of tone-locked responses further revealed that the context in which tones were presented modulated their evoked responses. These context effects were expressed as reduced N1 and N2 amplitudes for tones embedded in REG relative to RND sequences and an opposite effect (increased amplitudes for tones embedded in REG relative to RND) during the P2 response.

Overall, this pattern broadly aligns with previous findings reported by Hu et al. (24), who employed shorter silent gaps (200 ms; cycle duration 2,500 ms—approximately 50% shorter than in the present study). However, the effects observed here were weaker and emerged later in time. Whereas Hu et al. reported reliable regularity effects beginning in the second cycle, we observed such effects from the third cycle onward, consistent with weaker or less stable internal representations of the regularity (18).

The tone-evoked response profile also differed somewhat between studies. In Hu et al., tone-evoked responses were characterized primarily by two components (P1 and N1), with regularity effects (REG < RND) evident during the N1. In contrast, the responses observed here comprised a richer sequence of deflections. This difference likely reflects the longer inter-tone intervals used in the present study, which enabled the characterization of later evoked components, such as P2 and N2, that are not readily observable with shorter gaps. Differences between recording modalities (MEG in Hu et al., 2024, versus EEG here) may also have contributed to these discrepancies.

Notably, although reduced responses to REG relative to RND were observed during the N1 and N2 components, the opposite pattern emerged during the P2, indicating a transient reversal in the neural representation of regularity. This profile suggests a dynamic sequence of processing stages: an initial facilitatory effect for tones embedded in regular sequences (indexed by reduced N1 responses), followed by a reversal during the P2, and then a return to facilitation during the N2, where the strongest effect was observed (consistent with previous reports; e.g., (39). These findings indicate that regularity processing is governed by a hierarchy of processes operating at different stages of auditory analysis. Effects in the earlier components (N1/P2) emerged during the second regularity cycle, whereas the N2 effect— which also correlated with explicit short-term memory performance (see below)—appeared later, during the third cycle. This temporal dissociation suggests that the N2 may index a later stage of processing that is particularly dependent on mnemonic resources. Future work using more sensitive approaches, especially methods enabling reliable source localization, will be critical for clarifying the neural generators and functional significance of these effects.

We hypothesized that the substantial individual variability observed in the sequence evoked- and tone evoked-effects reflects differences in the capacity or efficiency of memory buffers required to bind individual tones over time and thereby detect repeating structure. PPM models provide a principled computational framework for linking predictive coding to memory, as predictions in these models are derived from stored representations of prior sequences across multiple context lengths (17,18). Crucially, the effective use of longer contexts, and thus the formation of precise predictions, depends on the fidelity and persistence of a memory buffer capable of maintaining sequential information over several seconds. When this buffer is limited or noisy, predictions must rely on shorter contexts, resulting in reduced precision and delayed detection of regularities (18). Individual differences in memory buffer capacity or persistence would therefore be expected to influence both the timing and strength of regularity detection, as well as the associated neural signatures. Within a PPM framework, such differences would manifest as variability in the maximum effective context length available to the observer, leading to weaker or later-emerging reductions in prediction error signals and attenuated sustained responses.

### Variability in explicit short-term memory capacity linked to variability in automatic EEG measures of sequence tracking

Consistent with previous work (28), we observed substantial inter-individual variability in performance on the explicit memory task, designed to probe the time- and capacity-limited processes responsible for temporarily retaining auditory information (55–57). The TP-COMP task uses stimuli closely matched to the REG/RND sequences employed during EEG recording, but requires participants to engage in an active, explicit delayed match-to-sample paradigm (58–61). Participants were asked to memorize a 500 ms tone pattern composed of 10 sequential 50 ms tones, retain it over a 2 s interval, and compare it to a subsequently presented probe. These rapid, arbitrary tone sequences minimize the possibility of rehearsal, allowing the task to probe low-level sensory representations rather than more explicitly structured or verbally mediated memory, such as in digit span tasks which have been commonly used in the literature to link memory processes and brain processing of sequences (e.g. (62); (63)). Neuroimaging and electrophysiological studies have identified core components of auditory short-term memory, including auditory cortex, as well as prefrontal (64–67) parietal regions (68–70) and the hippocampus (62,65,71).

Participants were naïve to the explicit memory task during EEG recording and completed it only at the end of the session. Although both tasks used the same tonal material, they differed markedly in their demands, allowing us to assess whether performance generalized across distinct forms of memory processing.

The central finding of this study is that individual variability in explicit memory performance was systematically related to both sustained and tone-locked neural responses during passive listening. Participants who performed well on the memory task exhibited larger sustained responses to REG relative to RND sequences, alongside reduced tone-locked responses - both signatures consistent with successful acquisition of the regularity. In contrast, participants with lower memory performance showed neither sustained response differences nor modulation of tone-evoked responses by sequence context.

In the tone-locked analysis, group differences between high- and low-memory performers were most pronounced in the N2 time window (250–400 ms post-onset). No significant group differences were observed in the N1 or P2 windows, suggesting that although these earlier components are sensitive to regularity (i.e., differ between REG and RND contexts), they are relatively insensitive to individual variability in memory performance. In contrast, the N2, often associated with contextual updating, mismatch evaluation, and the integration of incoming sensory input with learned regularities, appears to reflect processes that depend more directly on the quality or availability of memory representations. The selective modulation of the N2 therefore suggests that individual differences in memory capacity primarily impact later stages of processing involved in updating internal models of the auditory environment, rather than early sensory encoding.

Previous work has implicated the auditory N2 in memory-based prediction error processing. Todorovic and de Lange demonstrated that N2 responses are sensitive to violations of learned expectations even when sensory adaptation is controlled (39,72), suggesting a model-based rather than stimulus-driven origin. Bendixen et al. further emphasized the role of the N2 in comparing incoming input against stored representations of regularity, highlighting its dependence on the maintenance and integration of sequential information over time (3). Within a hierarchical predictive coding framework, Chennu et al. interpreted the N2 as a higher-level prediction error signal, reflecting violations of learned sequence structure rather than local acoustic features (73). Together, these findings support the interpretation of N2 modulation as a neural signature of context-dependent prediction error that depends on memory-based representations of temporal regularities.

### Sources of shared variability between short-term memory and automatic EEG responses

A key question concerns the nature of this shared variability. A simple explanation might be that it reflects differences in vigilance or attentional engagement, with participants who performed better on the memory task simply being more attentive during passive listening. Several observations argue against this interpretation. First, the EEG and memory tasks were temporally separated, and participants were unaware during EEG recording that a memory task would follow. Second, although high memory performers did exhibit larger onset responses, these effects were confined to early components and did not resemble canonical attentional enhancements, which typically extend to later evoked responses (74–76). Specifically, high performers showed stronger responses in the P1 time range, reliably linked to activity in primary and adjacent auditory cortex (e.g., Heschl’s gyrus and superior temporal plane), reflecting early cortical encoding of incoming sound (77), but not a generalized amplification of later components. This pattern is inconsistent with a generalized vigilance account and instead points to differences in early sensory encoding.

Another possibility is that the stronger onset responses observed in high memory performers reflect more efficient early sensory encoding, leading to the formation of more robust initial memory traces. Interestingly, although high performers exhibited differences in the early P1 and N1 components, these components were not the primary loci of memory-related effects in the main analyses. Instead, memory-related differences were most pronounced at later stages, indexed by the N2.

This pattern is consistent with a multi-stage account in which variability in early sensory encoding influences, but does not directly determine, subsequent processing stages. Stronger initial sensory representations may provide a more stable substrate for later processes involved in maintaining, integrating, or updating auditory context. In this view, individual differences in short-term memory performance may emerge not solely from differences in early encoding, but from how effectively early sensory information is transformed into sustained or contextually integrated representations that support predictive processing.

We therefore hypothesize that the observed effects reflect a shared memory buffer that supports both explicit memory performance and automatic regularity detection. Mechanistically, this buffer may involve hippocampal contributions—given the established role of the hippocampus in both sequence learning and explicit short-term memory (9,19,62,65,78)—or sustained activity within auditory cortical networks, which supports the maintenance of stimulus-specific representations over time (69,79–81). We refrained from source analysis due to the limited number of trials and the inherently noisy nature of EEG data. In future work source localization could help adjudicate between these possibilities.

In summary, our findings suggest that the memory processes engaged automatically during pattern scanning—and that play a canonical role in auditory predictive coding—are **not encapsulated**, but instead draw on resources shared with other cognitive functions. Listeners’ ability to detect regularities in the environment appears to depend on a common memory store, which may constitute a key source of inter-individual differences in predictive perception (e.g., (82)). Mechanistically, this shared substrate may link consciously accessible memory to automatic predictive processes either **directly**, through overlapping representational systems, or **indirectly**, via stronger sensory responses in auditory cortex that give rise to more robust and longer-lasting stimulus traces. Crucially, our results show that individual differences in automatic regularity detection can be predicted from a simple behavioural memory task, offering a tractable route for understanding variability in perceptual prediction. This link may extend to broader domains such as music perception, learning, and perceptual changes across the lifespan.

The present study focused on deterministic regularities. It therefore remains an open question whether the observed relationship between explicit memory performance and neural indices of regularity detection generalizes to stochastic or probabilistic structures, which impose different computational demands on memory and predictive mechanisms.

## Acknowledgments

This work was supported by a BBSRC project grant to MC. The funder had no role in study design, data collection, and analysis, decision to publish, or preparation of the manuscript.

## Data sharing

The data reported in this manuscript alongside related information will be available at DOI: XXX upon publication.

